## Supplemental data for "Genomic surveillance reveals that the dengue 2 virus lineage responsible for the 2023-2024 epidemic in the French Caribbean Islands is resistant to Mosnodenvir"

**Supplementary table 1a: Sequence information for the 98 sequences from the French Caribbean Island epidemic included in the analysis.** For each sequence, the Genbank accession number, strain name and collection date are provided.

| AN | Strain | Collection_date |
|---|---|---|
| **OR804009** | Mar-58 | 2023-04 |
| **PP326743** | 65244 | 2023-02-14 |
| **PP326744** | 65364 | 2023-02-18 |
| **PP326745** | 65866 | 2023-04-18 |
| **PP326746** | 65877 | 2023-04-19 |
| **PP326747** | 66202 | 2023-05-11 |
| **PP326748** | 66353 | 2023-04-04 |
| **PP326749** | 66354 | 2023-04-18 |
| **PP326750** | 66356 | 2023-04-24 |
| **PP326751** | 66357 | 2023-04-25 |
| **PP326752** | 66358 | 2023-04-25 |
| **PP326753** | 66361 | 2023-05-11 |
| **PP326754** | 66363 | 2023-05-12 |
| **PP326755** | 66364 | 2023-05-19 |
| **PP326756** | 66365 | 2023-05-22 |
| **PP326757** | 66366 | 2023-05-20 |
| **PP326758** | 66367 | 2023-05-19 |
| **PP326759** | 66368 | 2023-05-23 |
| **PP326760** | 66475 | 2023-06-06 |
| **PP326761** | 66482 | 2023-06-06 |
| **PP326762** | 66483 | 2023-06-07 |
| **PP326763** | 66484 | 2023-06-06 |
| **PP326764** | 66485 | 2023-06-07 |
| **PP326765** | 66593 | 2023-06-09 |
| **PP326766** | 66594 | 2023-06-13 |
| **PP326767** | 66595 | 2023-06-12 |
| **PP326768** | 66596 | 2023-06-12 |
| **PP326769** | 66597 | 2023-06-13 |
| **PP326770** | 66897 | 2023-06-21 |
| **PP326771** | 66898 | 2023-06-22 |

| | | |
|---|---|---|
| **PP326772** | 66901 | 2023-06-27 |
| **PP326773** | 66902 | 2023-06-26 |
| **PP326774** | 66904 | 2023-06-27 |
| **PP326775** | 66905 | 2023-06-27 |
| **PP326776** | 66909 | 2023-06-29 |
| **PP326777** | 67008 | 2023-07-03 |
| **PP326778** | 67187 | 2023-07-03 |
| **PP326779** | 67188 | 2023-07-03 |
| **PP326780** | 67189 | 2023-07-05 |
| **PP326781** | 67190 | 2023-07-05 |
| **PP326782** | 67192 | 2023-07-06 |
| **PP326783** | 67198 | 2023-07-07 |
| **PP326784** | 67201 | 2023-07-11 |
| **PP326785** | 67561 | 2023-07-26 |
| **PP326786** | 67618 | 2023-08-02 |
| **PP326787** | 68064 | 2023-08-09 |
| **PP326788** | 68252 | 2023-08-23 |
| **PP326789** | 68637 | 2023-08-16 |
| **PP326790** | 68638 | 2023-08-19 |
| **PP326791** | 68641 | 2023-08-17 |
| **PP326792** | 68645 | 2023-08-21 |
| **PP326793** | 68649 | 2023-08-17 |
| **PP326794** | 68650 | 2023-08-23 |
| **PP326795** | 68651 | 2023-08-19 |
| **PP326796** | 68652 | 2023-08-18 |
| **PP326797** | 68925 | 2023-08-23 |
| **PP326798** | 68932 | 2023-08-24 |
| **PP326799** | 69057 | 2023-09-03 |
| **PP326800** | 69212 | 2023-08-29 |
| **PP326801** | 69225 | 2023-08-30 |
| **PP326802** | 69227 | 2023-09-06 |
| **PP326803** | 69709 | 2023-09-05 |
| **PP326804** | 69710 | 2023-09-05 |
| **PP326805** | 69711 | 2023-09-07 |

| | | |
|---|---|---|
| **PP326806** | 69712 | 2023-09-07 |
| **PP326807** | 69714 | 2023-09-09 |
| **PP326808** | 69716 | 2023-09-11 |
| **PP326809** | 69717 | 2023-09-12 |
| **PP326810** | 69718 | 2023-09-12 |
| **PP326811** | 69947 | 2023-09-18 |
| **PP326812** | 69948 | 2023-09-18 |
| **PP326813** | 70053 | 2023-09-26 |
| **PP326814** | 70621 | 2023-10-12 |
| **PP326815** | 70686 | 2023-10-12 |
| **PP326816** | 70694 | 2023-08-29 |
| **PP326817** | 70923 | 2023-10-21 |
| **PP326818** | 70972 | 2023-10-24 |
| **PP326819** | 71023 | 2023-10-05 |
| **PP326820** | 71024 | 2023-10-06 |
| **PP326821** | 71027 | 2023-10-05 |
| **PP326822** | 71050 | 2023-08-07 |
| **PP326823** | 71056 | 2023-09-12 |
| **PP326824** | 71057 | 2023-08-24 |
| **PP326825** | 71333 | 2023-11-01 |
| **PP326826** | 71492 | 2023-10-24 |
| **PP326827** | 71874 | 2023-11-20 |
| **PP326828** | 72475 | 2023-12-13 |
| **PP326829** | 72487 | 2023-12-14 |
| **PP326830** | 72527 | 2023-12-11 |
| **PP326831** | 72536 | 2023-12-19 |
| **PP326832** | 72674 | 2023-12-28 |
| **PP326833** | 72728 | 2024-01-03 |
| **PP326834** | AE-BV-B4 | 2023-10-16 |
| **PP331235** | 66906 | 2023-06-27 |
| **PP331236** | 68639 | 2023-08-17 |
| **PP335480** | 72533 | 2023-12-14 |
| **PP335483** | 66362 | 2023-05-12 |
| **PP335484** | 68628 | 2023-08-22 |

**Supplementary table 1b: Prevalence among sequences from the French Caribbean Islands for described JNJ A07 resistance mutations.** The percentage of sequences exhibiting the mutation is provided for each mutation.

| NS4B mutation | Prevalence in French Caribbean Islands sequences (%) |
|---|---|
| **F47Y** | 0 |
| **S85L** | 0 |
| **V91A** | 100 |
| **L94F** | 0 |
| **P104S** | 0 |
| **T108I** | 0 |
| **A137T** | 0 |
| **T216N** | 0 |
| **T216P** | 0 |

**Supplementary table 2:** Provided as a supplementary file

**Supplementary table 3: DENV-2 V91A-carrying sequences information.** For each DENV-2 sequence carrying mutation V91A, the Genbank accession number, geographic origin, collection year, and genotype are provided.

| Accession number | Geographic origin | Year | Genotype |
|---|---|---|---|
| PP269912 | Viet Nam | 2023 | Cosmopolitan |
| PP269905 | Viet Nam | 2022 | Cosmopolitan |
| PP269889 | Viet Nam | 2019 | Asian I |
| PP269883 | Viet Nam | 2019 | Asian I |
| OR821962 | USA | 2023 | Cosmopolitan |
| OR771147 | USA | 2023 | Cosmopolitan |
| OQ028215 | Viet Nam | 2019 | Asian I |
| OQ028224 | Viet Nam | 2019 | Asian I |
| MW512438 | Singapore | 2015 | Cosmopolitan |
| MW512459 | Singapore | 2016 | Cosmopolitan |
| MW946255 | Thailand | 2005 | Asian I |
| MW946309 | Thailand | 2004 | Asian I |
| MW946352 | Thailand | 2007 | Asian I |
| MW946353 | Thailand | 2004 | Asian I |

| | | | |
|---|---|---|---|
| MW946449 | Thailand | 2014 | Asian I |
| MW946517 | Thailand | 2005 | Asian I |
| MW946531 | Thailand | 2010 | Asian I |
| MW946595 | Thailand | 2005 | Asian I |
| MN448838 | Thailand | 2010 | Asian I |
| MN448905 | Thailand | 2009 | Asian I |
| MN448907 | Thailand | 2009 | Asian I |
| MG779202 | Kenya | 2017 | Cosmopolitan |
| JN819419 | Brazil | 2000 | Asian American |
| HQ999999 | Guatemala | 2009 | Asian American |
| FJ810410 | Thailand | 2001 | Asian I |
| FJ687434 | Thailand | 2001 | Asian I |
| FJ687435 | Thailand | 2001 | Asian I |
| FJ687436 | Thailand | 2001 | Asian I |
| FJ687444 | Thailand | 2001 | Asian I |
| FJ687445 | Thailand | 2001 | Asian I |
| FJ687446 | Thailand | 2001 | Asian I |
| FJ744719 | Thailand | 2001 | Asian I |
| FJ744720 | Thailand | 2001 | Asian I |
| FJ744721 | Thailand | 2001 | Asian I |
| FJ744722 | Thailand | 2001 | Asian I |
| FJ744723 | Thailand | 2001 | Asian I |
| FJ639828 | Thailand | 2001 | Asian I |
| FJ639829 | Thailand | 2001 | Asian I |
| FJ639830 | Thailand | 2001 | Asian I |

**Supplementary table 4: DENV-3 V91A-carrying sequences information.** For each DENV-3 sequence carrying mutation V91A, the Genbank accession number, geographic origin, collection year, and genotype are provided.

| Accession number | Geographic origin | Year | Genotype |
|---|---|---|---|
| EU660410 | Viet Nam | 2006 | II |
| MZ008477 | Nicaragua | 2013 | III |
| MZ008475 | Nicaragua | 2013 | III |
| MW946935 | Thailand | 1997 | II |
| MW946852 | Thailand | 2006 | II |
| MW946829 | Thailand | 2005 | II |
| KU509284 | Thailand | 2008 | II |
| KF973486 | Nicaragua | 2012 | III |
| KF921927 | Nicaragua | 2010 | III |
| OQ821525 | Cuba | 2022 | III |

**Supplementary Figure 1: DENV-2 V91A-carrying sequences genotypes.** Maximum-Likelihood phylogeny of all V91A-carrying DENV2 sequences with a set of reference sequences representative of DENV-2 genotypes Phylogenetic inference was performed using IQTREE2 under the best substitution model identified by ModelFinder with ultrafast bootstrap approximation (1000 replicates). The tree was rooted using the highly divergent strain QML22 (KX274130). Nodes with boostrap support above 95 are highlighted with a black circle.

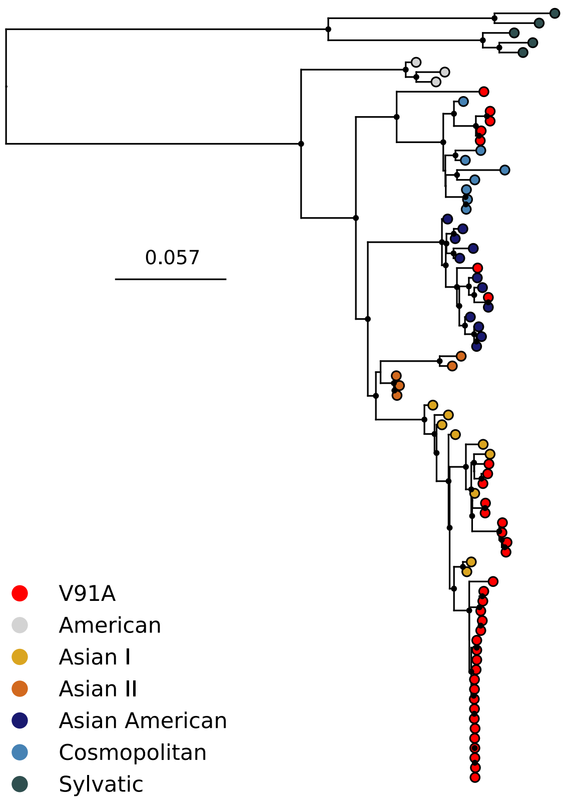

**Supplementary Figure 2: DENV-3 V91A-carrying sequences genotypes.** Maximum-Likelihood phylogeny of all V91A-carrying DENV2 sequences with a set of reference sequences representative of DENV-3 genotypes Phylogenetic inference was performed using IQTREE2 under the best substitution model identified by ModelFinder with ultrafast bootstrap approximation (1000 replicates). The tree was midpoint rooted. Nodes with boostrap support above 95 are highlighted with a black circle.

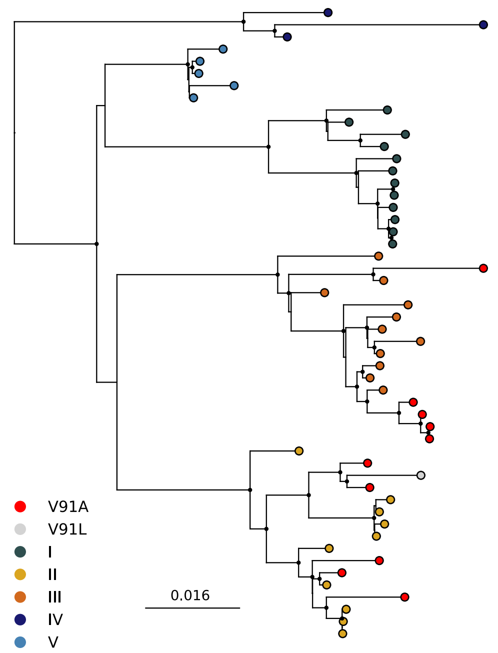
